# Assessing the efficacy of therapeutically promising combination of polymyxin B and triclosan against colistin-resistant *Klebsiella pneumoniae*

**DOI:** 10.1101/2024.12.17.628820

**Authors:** Soumya Biswas, Pushney Mahapatra, Soujanya Ghosh, Saumya Darshana Patra, Gaurav Verma, Namrata Misra, Gajraj Singh Kushwaha, Mrutyunjay Suar

## Abstract

Antimicrobial resistance (AMR) is one of the greatest public health challenges of the current time, and it is primarily associated with Gram-negative bacterial infections. Among these infections, *Klebsiella pneumoniae* is the most notorious bacterial pathogen in the context of AMR dissemination. In this dire circumstance, clinicians explored old regimes such as colistin to treat these multidrug-resistant infections; however, unfortunately, the resistance towards these last-resort drugs is also emerging rapidly. In this study, we examined the occurrence of colistin resistance in *K. pneumoniae* concerning its mechanism of resistance, chronology, and geographical distribution. We found that resistance towards these last-resort drugs, colistin and polymyxin B, is emerging rapidly, with evolving multiple mechanisms of resistance. While efforts continue to develop new antibacterial drugs, a combination of approved antibacterial drugs may be one of the most suitable strategies to fight against AMR in the current time-ticking situation. In this series, we evaluated the efficacy of a therapeutically viable combination of polymyxin B and triclosan using a set of standard antibacterial assays, used for *in vitro* preclinical efficacy. We found that the combination is highly effective in colistin-resistant clinical strains of *K. pneumoniae,* as triclosan sensitizes the pathogen towards polymyxin B. Furthermore, the results of additional complementary experiments, such as time-kill kinetics, biofilm inhibition, frequency of resistance, cell viability, and cytotoxicity assays, show very encouraging results for the feasibility and validation of this combination. Finally, we examined the presence of mutations in field isolates of *K. pneumoniae* FabI enzyme, as it has been hypothesized previously that triclosan may cause mutation in its binding target, FabI. Interestingly, we could not detect any mutation in the triclosan binding site of FabI in the genome of field isolates. Altogether, this study provides an overview of the current situation on colistin resistance, a promising solution for colistin resistance comprising a combination of two already being used antibacterial ingredients based on *in vitro* efficacy studies.

## 1. Introduction

*Klebsiella pneumoniae*, infamous for multidrug resistance, is a leading cause of nosocomial infections, including pneumonia, urinary tract infections, sepsis, and meningitis (Podschun and Ullmann, 1998). In the recent few decades, *K. pneumoniae* has been identified by World Health Organization (WHO) as a major concern for antimicrobial resistance (AMR) due to its high rate of resistance to commonly used antibiotics, such as penicillin, quinolones, and cephalosporins (Chen et al., 2014; Henson et al., 2017; Rawat and Nair, 2010). The emergence of multidrug-resistant (MDR) strains of *K. pneumoniae*, including extended-spectrum beta-lactamase-producing (ESBL), carbapenem-resistant *K. pneumoniae* (CRKP), poses an urgent public health challenge (Gupta et al., 2011; Pitout and Laupland, 2008; Van Duin et al., 2013).

The alarming rise of MDR Gram-negative pathogens resistant to aminoglycosides, fluoroquinolones, and β-lactams, including carbapenems, has driven a consideration of old regimens such as colistin and polymyxin B as a “last-resort” therapeutic option for critically ill patients (Bialvaei and Samadi Kafil, 2015). Unfortunately, the widespread use of colistin and polymyxin B has driven the emergence of resistance in *K. pneumoniae*. The first reported case of colistin-resistant *K. pneumoniae* was in May 2004 (Antoniadou et al., 2007) though the precise molecular mechanism of resistance was not fully understood at that time. Later studies revealed that colistin resistance in Gram-negative bacterial pathogens is associated with multiple mechanisms, including mutations in *mgrB* gene, PmrAB and PhoPQ operons, and crrAB two-component system (Aghapour et al., 2019). A breakthrough occurred in 2016 with the discovery of a plasmid-mediated colistin resistance gene, mobile colistin resistance-1 (*mcr-1*) (Liu et al., 2016). Within a year, *mcr-1* gene was detected across the globe, including Asia, Europe, North America, and South America (Skov and Monnet, 2016; Zhu et al., 2021).

Although new classes of antibacterial drugs are in the pipeline, approval of these new drugs against novel targets in Gram-negative pathogens remains a significant challenge (Andrei et al., 2019). Consequently, exploring novel approaches, such as a combination of approved compounds, shows promising results in combating bacterial AMR and enhancing therapeutic strategies for treating MDR infections (Tyers and Wright, 2019; Worthington and Melander, 2013). Recently, a combination of antibiotics with non-antibiotic compounds has emerged as a highly promising strategy for addressing AMR (Faruqe, 2024; Koh Jing Jie et al., 2022; Xiao et al., 2023), and among them, fatty acid synthesis inhibiting compounds have shown promising results against colistin resistance in *K. pneumoniae* (Carfrae et al., 2023). Triclosan [5-chloro-2-(2,4-dichlorophenoxy) phenol: TCL] primarily targets the bacterial fatty acid synthesis (FAS II) pathway by inhibiting the activity of enoyl acyl-carrier protein reductase (ENR or FabI) (Levy et al., 1999; McMurry et al., 1998).

In this study, we found that the prevalence of colistin resistance in *K. pneumoniae* strains has risen rapidly from 2009 to 2023 by analysing over 36,000 whole-genome sequences. Similarly, colistin-resistant gene, *mcr-1* occurrence has also increased since its detection in 2016, while *mcr-1*-positive cases remain fewer compared to colistin-resistant cases without *mcr-1*. Checkerboard assays on colistin-resistant clinical strains of *K. pneumoniae* showed a strong synergy, reducing polymyxin B MICs up to 16-fold (*mcr-1*-positive) and 128-fold (*mcr-1*-negative), and triclosan MICs by over 200-fold. The combination showed an excellent average fractional inhibitory concentration index of 0.0759 (*mcr-1*-positive) and 0.015 (*mcr-1*-negative), and Synergy Finder analysis reported a zero-interaction potency score of 27.78. Furthermore, time-kill assays indicated complete bacterial eradication within 48 hours at sub-MIC concentrations. Biofilm formation was fully inhibited, and cytotoxicity testing on RAW 264.7 cells showed minimal toxicity. Finally, variant-calling analysis of the 36,000 isolates revealed that no significant mutations were observed in the FabI binding site despite the potential of triclosan to induce resistance.

## 2. Results

### 2.1. *mcr-1* gene is not solely responsible for colistin resistance in *Klebsiella pneumoniae*

The analysis of *K. pneumoniae* whole-genome data from 2009 to 2023 showed a significant and alarming increase in colistin resistance over time (Fig. 1). Resistance rates, which ranged between 4% and 8% in earlier years, escalated sharply to over 12% by 2023, indicating an upward trend that poses a serious challenge to the efficacy of colistin as a last-resort treatment. The plasmid-mediated *mcr-1* gene, a key contributor to colistin resistance, was first detected in 2016 in isolated cases. Since then, its prevalence has gradually increased, peaking in 2023 at less than 0.0004%. However, the significant disparity between the total number of colistin-resistant cases (2597) and *mcr-1* positive cases (88) strongly suggests that alternative resistance mechanisms, such as chromosomal mutations or non-*mcr-1* plasmid-mediated pathways, play a predominant role in driving colistin resistance in *K. pneumoniae*.

**Figure 1:**
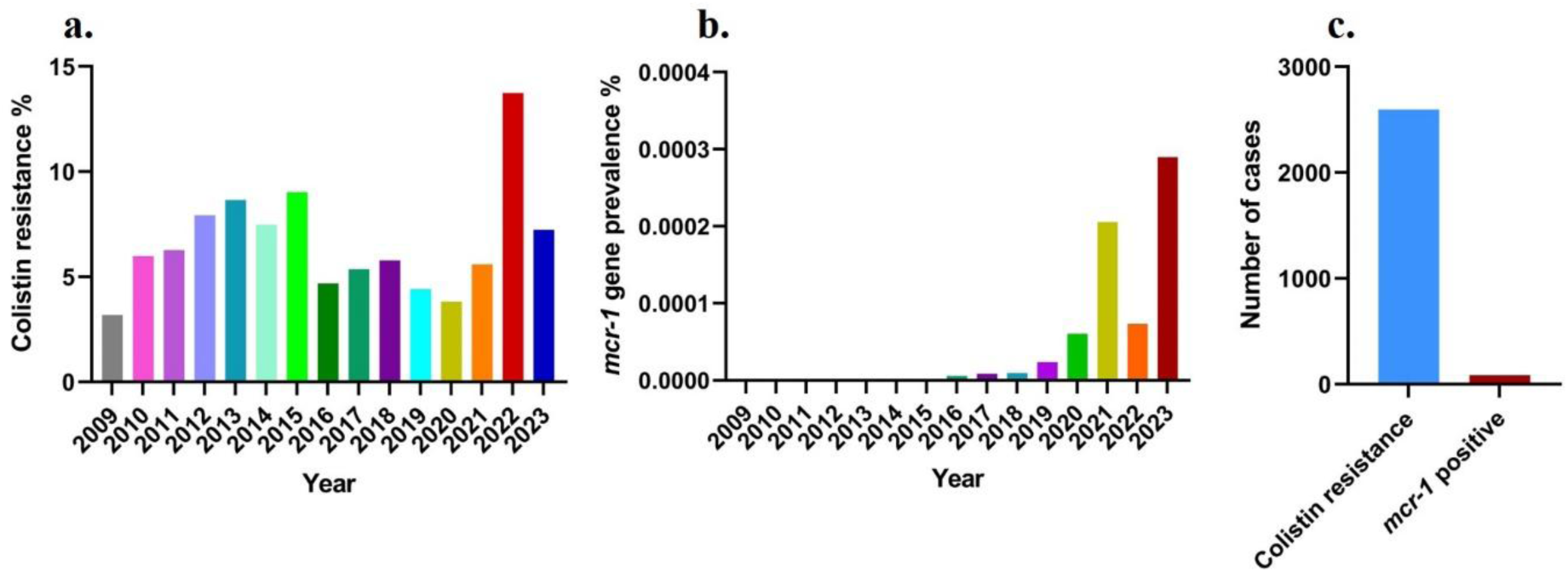
The prevalence of colistin resistance and the emergence of the *mcr-1* gene in *K. pneumoniae* have increased significantly between 2009 and 2023. **(a.)** The proportion of *K. pneumoniae* isolates resistant to colistin rose from about 5% in 2009 to over 15% by 2023. **(b.)** The *mcr-1* gene, a primary resistance gene in colistin-resistant pathogens, was detected in 2016, and its prevalence climbed to over 0.003% by 2023. **(c.)** The total number of colistin-resistant cases grew from a few hundred in 2009 to more than 2,500 in 2023. Among these, *mcr-1*-positive cases increased from nearly zero to 88 by 2023.

### 2.2. Global prevalence of colistin resistance in *Klebsiella pneumoniae*

The map illustrates the global distribution of *K. pneumoniae* sequences exhibiting colistin resistance for the period 2009 to 2023 (Fig. 2). Europe reports the highest number of cases, with 948 sequences, indicating a significant burden of colistin-resistant *K. pneumoniae* in this region. North America follows with 353 cases, reflecting a substantial prevalence, while Asia reports 293 cases, highlighting its role as a significant region for resistance emergence. Africa accounts for 100 cases, and South America and Australia report lower numbers, with 42 and 27 cases, respectively. These findings emphasize the global dissemination of colistin resistance in *K. pneumoniae*, with particularly high prevalence in Europe and North America. This data emphasises the critical need for enhanced surveillance systems and region-specific strategies to mitigate the spread of resistance and address this pressing public health challenge.

**Figure 2:**
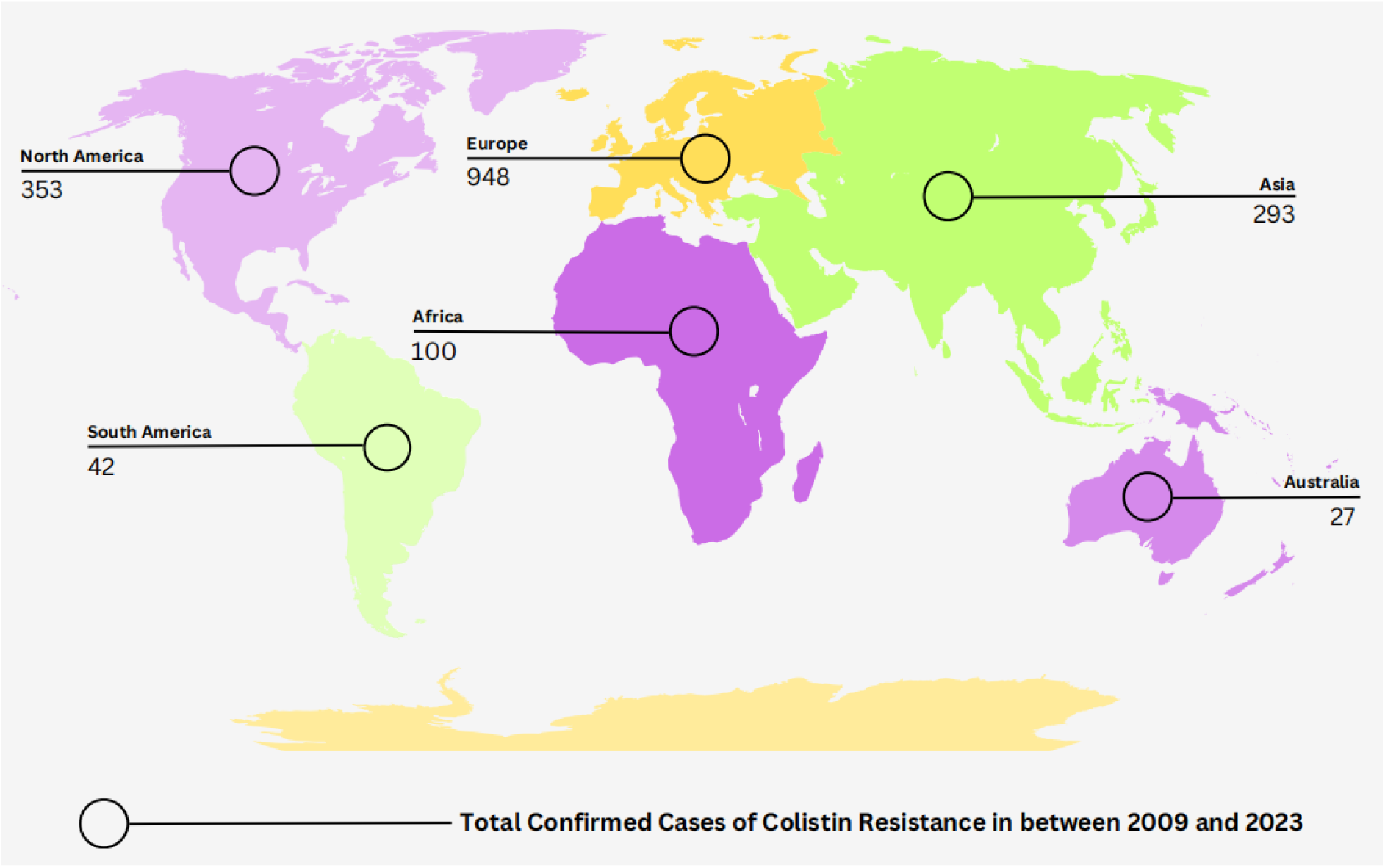
Global distribution of *K. pneumoniae* sequences exhibiting colistin resistance. The map highlights the regional variability in resistance cases, with Europe showing the highest number of confirmed sequences (948), followed by North America (353) and Asia (293). Africa reports 100 cases, while South America and Australia have significantly fewer cases, with 42 and 27, respectively.

### 2.3. Triclosan synergistically potentiates polymyxin B activity in colistin-resistant *Klebsiella pneumoniae*

First, six colistin-resistant clinical strains were isolated and identified using a susceptibility assay, followed by PCR amplification of *mcr-1* gene (three *mcr-1* positive and three *mcr-1* negative). In all cases, the combination assays resulted in a marked reduction in the MICs of both agents compared to monotherapy (Figure 3, a-f). The calculated fractional inhibitory concentration indices (FICᵢ) ranged from 0.0095 to 0.0925, indicating strong synergy in all strains (FICᵢ < 0.5) (Figure 3, g).

**Figure 3:**
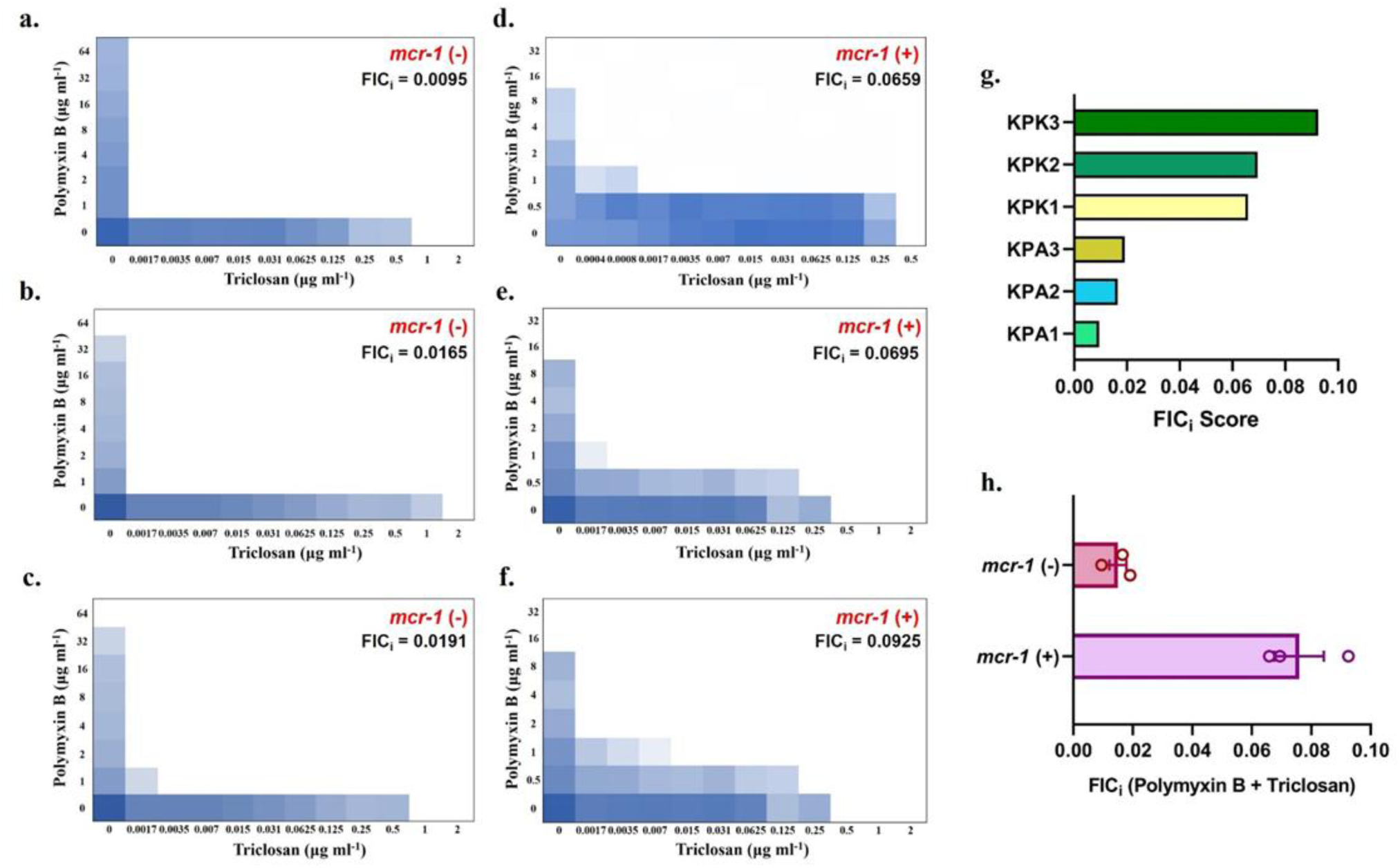
Synergistic activity of triclosan and polymyxin B against *K. pneumoniae*, including *mcr-1*-positive colistin-resistant strains. **a.-c**. Checkerboard broth microdilution assays showing the combinatorial effects of polymyxin B and triclosan on *mcr-1*-negative, colistin-resistant *K. pneumoniae* clinical isolates, KPA1 **(a.)**, KPA2 **(b.)**, and KPA3 **(c.). d–f.** Synergistic efficacy between triclosan and polymyxin B in *mcr-1*-positive *K. pneumoniae* clinical isolates, KPK1 **(d.)**, KPK2 **(e.)**, and KPK3 **(f.)**. The degree of inhibition is visualized using a heat map, with darker shades indicating higher growth inhibition. (**g.)** Quantitative comparison of FICᵢ values across six clinical *K. pneumoniae* isolates. A horizontal bar graph illustrates the FICᵢ values for each strain: KPA1–KPA3 (*mcr-1*-negative) and KPK1–KPK3 (*mcr-1*-positive). Each FICi score for individual strains was determined experimentally with at least three independent replicates to ensure reproducibility. (**h.)** Comparative analysis of FICᵢ values for *mcr-1*-positive and *mcr-1-*negative *K. pneumoniae* isolates treated with a combination of polymyxin B and triclosan. The bar plot displays the mean FICᵢ values for each group, with individual data points representing biological replicates (n = 3 per strain). Error bars indicate the full range of FICᵢ values observed across replicates. Despite a statistically significant difference in mean FICᵢ values between the two groups (p < 0.05, unpaired t-test), both remain well within the synergistic range (FICᵢ < 0.5).

In *mcr-1*-positive strains, polymyxin B MICs were reduced from 16 µg/mL to 1 µg/mL (93.75% reduction), while triclosan MICs dropped from 0.5 µg/mL to 0.0017-0.015 µg/mL, representing up to 99.66% reduction (Table 1). In *mcr-1*-negative strains, the MICs of polymyxin B declined from 64-128 µg/mL to 1 µg/mL (98.43 to 99.21% reduction), and triclosan MICs were reduced from 1-2 µg/mL to 0.0017-0.0035 µg/mL (up to 99.91% reduction). Although *mcr-1*-positive strains exhibited slightly higher FICᵢ values and a lower degree of MIC reduction compared to *mcr-1*-negative strains (Figure 3, h), the difference was statistically significant, as determined by an unpaired two-tailed t-test (p < 0.05). Despite this reduced magnitude of synergy, the combination still consistently fell within the synergistic range (FICᵢ < 0.5), indicating therapeutic potential even in resistant strains. Based on this observation, we used the *mcr-1*-positive strain (KPK1) for further experiments, as it represents a more clinically challenging and relevant population due to its colistin resistance.

**Table 1:** Synergistic antibacterial activity of polymyxin B combined with triclosan against *mcr-1* positive and *mcr-1* negative clinical strains of *K. pneumoniae*.

| Strains | *mcr-I* | Drug Combination | Polymyxin B |  | Percentage of reduction | Triclosan |  | Percentage of reduction | FIC index | Effect |
| --- | --- | --- | --- | --- | --- | --- | --- | --- | --- | --- |
|  |  |  | MIC alone | MIC in Combination |  | MIC alone | MIC in Combination |  |  |  |
| KPK1 | + | PMB/TCS | 16 | 1 | 93.75% | 0.5 | 0.0017 | 99.66% | 0.0659 | Synergistic |
| KPK2 | + | PMB/TCS | 16 | 1 | 93.75% | 0.5 | 0.0035 | 99.30% | 0.0695 | Synergistic |
| KPK3 | + | PMB/TCS | 16 | 1 | 93.75% | 0.5 | 0.015 | 97.00% | 0.0925 | Synergistic |
| KPA1 | - | PMB/TCS | 128 | 1 | 99.21% | 1 | 0.0017 | 99.83% | 0.0095 | Synergistic |
| KPA2 | - | PMB/TCS | 64 | 1 | 98.43% | 2 | 0.0017 | 99.91% | 0.0165 | Synergistic |
| KPA3 | - | PMB/TCS | 64 | 1 | 98.43% | 1 | 0.0035 | 99.65% | 0.0191 | Synergistic |

**Table 2:** Single-step mutation frequencies of *Klebsiella pneumoniae* under polymyxin B, triclosan, and combination

| Drug | MIC ($\mu\text{g/mL}$) | Drug Conc. ($\mu\text{g/mL}$) | Inoculum (CFU/mL) | Single-Step Mutation frequency |
| --- | --- | --- | --- | --- |
| Polymyxin B | 16 | 64 | $2.8 \times 10^9$ | $4.6 \times 10^{-9}$ |
| | | 128 | | $1.07 \times 10^{-9}$ |
| Triclosan | 0.5 | 2 | $2.6 \times 10^9$ | $< 2.6 \times 10^{-9}$ |
| | | 4 | | $< 2.6 \times 10^{-9}$ |
| PMB + TCS | — | 16 + 0.5 | $2.7 \times 10^9$ | $< 2.7 \times 10^{-9}$ |

Subsequently, the synergy of KPK1 was further quantified using the SynergyFinder application (version 3.0) (Ianevski et al., 2022), which generated a ZIP synergy score of 27.78, validating the pronounced synergy between the two antibiotics. Visual analyses, including a 2D synergy map and 3D δ-score plots, highlighted regions of optimal synergy across various concentration combinations, while the inhibition heatmap confirmed near-complete microbial inhibition (100%) in specific combinations (Fig. 4), which then guided the dose selection for time-kill assays and other downstream experiments.

**Figure 4:**
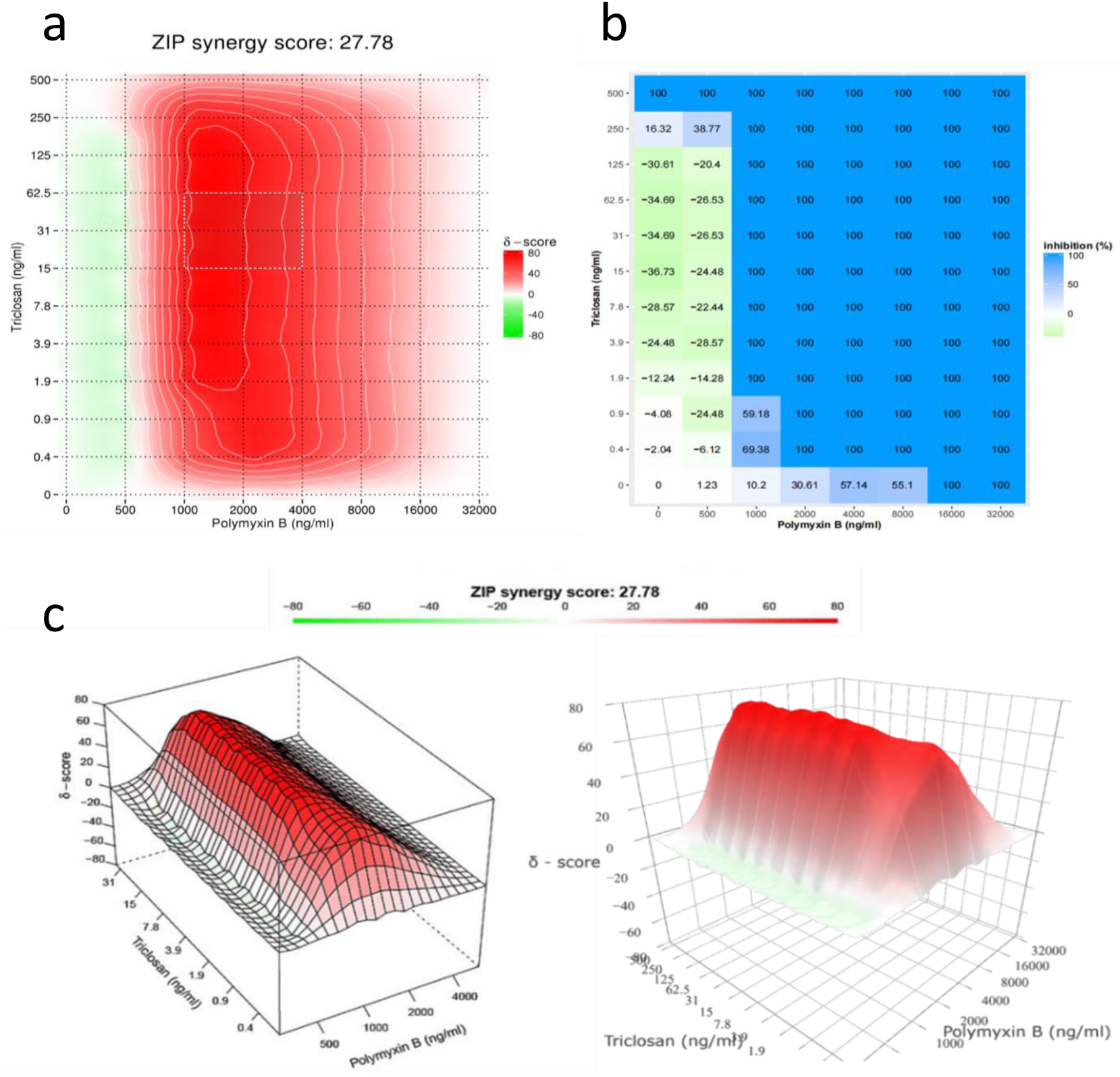
Synergistic interaction between polymyxin B and triclosan against KPK1 was analyzed using checkerboard broth microdilution and visualized through synergy mapping. Regions of optimal synergy are highlighted across concentration ranges. **(a.)** 2D synergy map displaying the δ-scores of triclosan and polymyxin B combinations. Red regions indicate strong synergy, with a ZIP synergy score of 27.78, confirming enhanced antimicrobial activity, and the map highlights regions where optimal synergy is achieved across various concentrations. **(b.)** Heatmap showing inhibition percentages across various concentration combinations. Maximum inhibition (100%) is observed in specific drug pair concentrations. **(c.)** 3D visualization of the δ-scores, providing a topographical view of synergy intensity, with clear peaks representing optimal synergistic combinations.

### 2.4. Time-dependent eradication of colistin-resistant *Klebsiella pneumoniae*

The time-kill kinetics assay for colistin-resistant *K. pneumoniae* (KPK1) showed that neither triclosan (TCS) nor polymyxin B (PMB) at sub-MIC concentrations (1/8 MIC) exhibited significant antibacterial activity when used alone (Fig. 5). Triclosan treatment (0.0625 µg/mL) allowed bacterial growth to progress similarly to the growth control, with counts reaching approximately 10^11^ CFU/mL by 24 hours. Similarly, polymyxin B treatment (2 µg/mL) showed minimal effect, with bacterial counts remaining close to those of the growth control throughout the experiment. In contrast, the combination of triclosan and polymyxin B at 1/8 MIC each displayed a potent synergistic effect. A significant reduction in bacterial counts was observed by 8 hours, with levels decreasing to approximately 10^3^ CFU/mL by 24 hours. Notably, after 48 hours of combination treatment, no bacterial colonies were observed, indicating complete eradication of the bacterial population.

**Figure 5:**
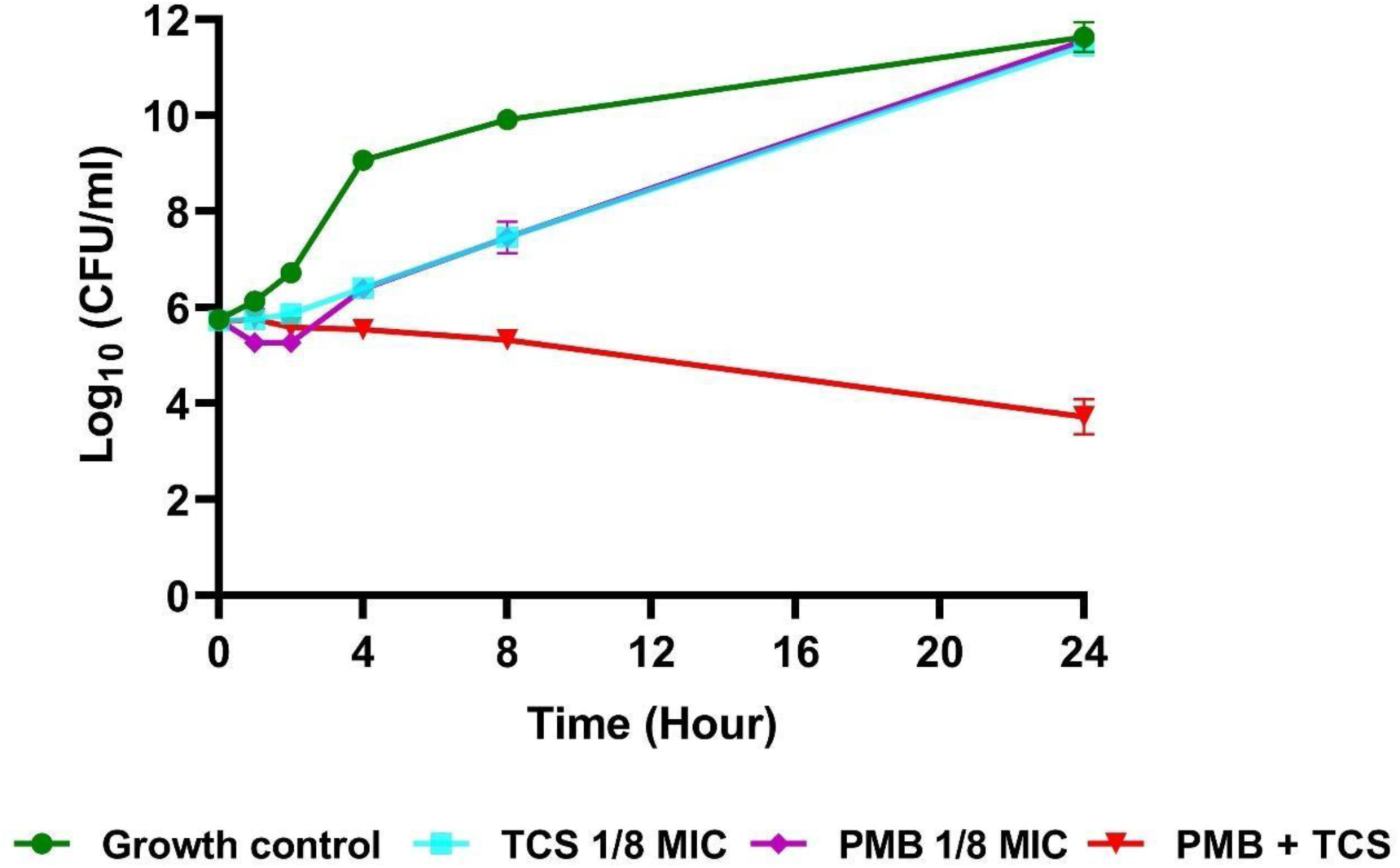
The time-kill assay shows the antimicrobial effects of triclosan, polymyxin B, and their combination at sub-MIC concentrations (1/8 MIC each) against colistin-resistant *K. pneumoniae*. The x-axis represents time (0−24 hours), and the y-axis indicates bacterial counts (Log_10_ CFU/mL). The growth control (green line) and triclosan alone (cyan line) both exhibit exponential bacterial growth, reaching approximately 10^11^ CFU/mL by 24 hours, while polymyxin B alone (magenta line) has a minimal inhibitory effect, with bacterial counts close to the control. In contrast, the combination of triclosan and polymyxin B (red line) shows a strong synergistic effect, significantly reducing bacterial counts by 8 hours and achieving approximately 10^3^ CFU/mL by 24 hours. The absence of colonies after 48 hours of combination treatment, although not shown in the graph, confirms complete bacterial eradication. Data represent the mean ± standard deviation from triplicate samples.

### 2.5. Combination prevents biofilm formation

We evaluated the effects of sub-minimal inhibitory concentrations (sub-MICs) of polymyxin B (PMB) (2 µg/mL, 1/8 MIC) and triclosan (TCS) (0.0625 µg/mL, 1/8 MIC), individually and in combination, on biofilm formation. The crystal violet staining assay demonstrated that the PMB + TCS combination significantly inhibited biofilm formation compared to the untreated growth control (GC) (Fig. 6). Individual treatments with PMB or TCS produced only minor reductions in biofilm formation, as their absorbance values were slightly lower than the GC but not significantly different. Visual inspection of the stained 96-well plate supported these findings, with wells treated with PMB or TCS alone showing staining levels similar to the GC, indicating minimal biofilm disruption. In contrast, the PMB + TCS combination-treated wells exhibited almost no staining, reflecting an almost complete inhibition of biofilm formation. A repeated measures ANOVA was also applied, which revealed a significant effect of treatment (F(1.139, 2.277) = 72.88, P = 0.0089). Due to a violation of the sphericity assumption (ε = 0.3795), the Geisser-Greenhouse correction was applied. There was no significant variation among triplicates of each group, indicating consistent responses across individuals.

**Figure 6:**
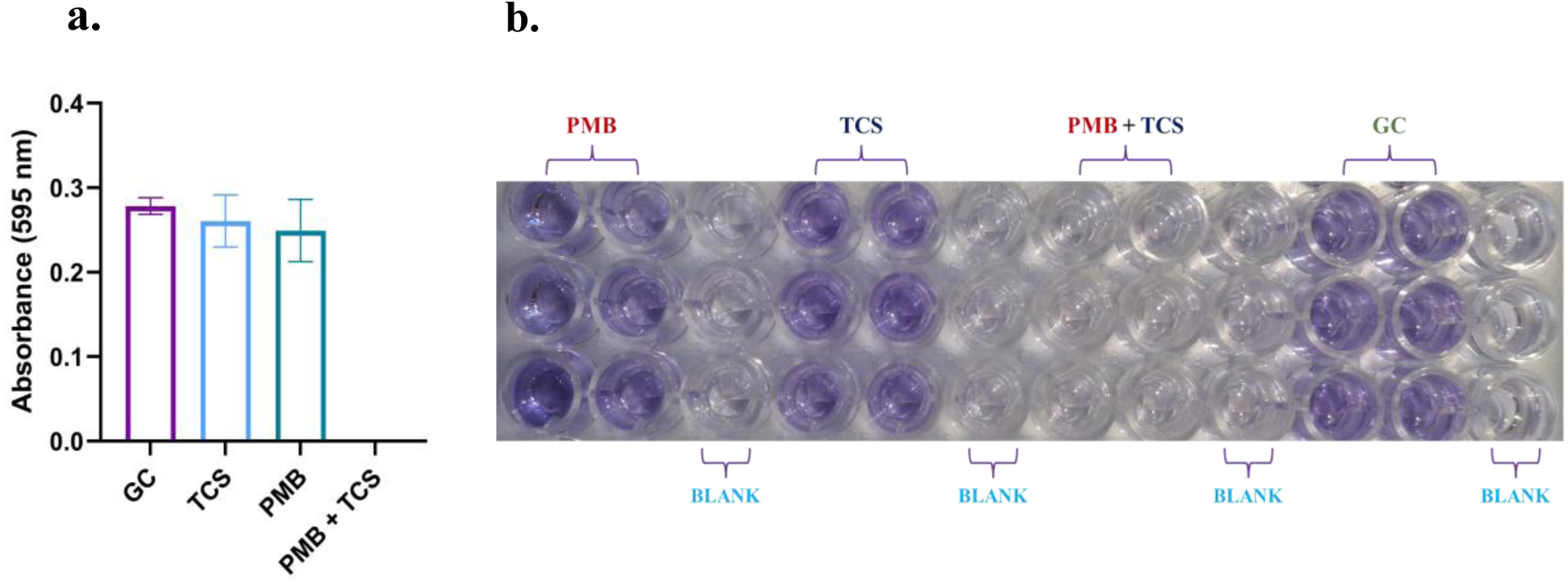
Crystal violet staining was used to quantify biofilm formation by *K. pneumoniae* (KPK1) in the presence of sub-minimal inhibitory concentrations (1/8 MIC) of polymyxin B (PMB, 2 µg/mL) and triclosan (TCS, 0.0625 µg/mL), individually and in combination. **(a.)** Absorbance data at 595 nm demonstrate that the PMB + TCS combination completely inhibited biofilm formation, while individual treatments with PMB or TCS resulted in biofilm levels only slightly lower than the growth control (GC). Data are presented as mean ± standard deviation from three independent experiments. Statistical analysis was performed using repeated measures ANOVA with Geisser-Greenhouse correction (F(1.139, 2.277) = 72.88, *P* = 0.0089), indicating a significant treatment effect. **(b.)** Visual confirmation of biofilm biomass via crystal violet staining in a 96-well plate. Wells treated with PMB + TCS show no visible staining, indicating no biofilm formation, while those treated with PMB or TCS alone exhibit staining similar to the GC. Blank wells serve as negative controls.

### 2.6. *Klebsiella pneumoniae* showed insignificant resistance evolution against the combination

The mutation frequencies in *K. pneumoniae* were analysed under selective pressure from polymyxin B, triclosan, and their combination at varying concentrations. Polymyxin B exhibited mutation frequencies of 4.6 × 10^-9^ and 1.07 × 10^-9^ at 4× MIC and 8× MIC, respectively, indicating a significant reduction in resistant mutants at higher drug concentrations. For triclosan, no resistant colonies were observed at 4× and 8× MIC range, with mutation frequency below the detectable threshold relative to the inoculum size. Similarly, the combination treatment did not result in detectable colonies at the MIC level, signifying an absence of mutations under these conditions. Furthermore, since no growth was observed at 4× MIC of triclosan, it suggests that resistance against triclosan is also unlikely to develop.

### 2.7. Combination potentiates membrane disruption

The flow cytometry analysis revealed the impact of sub-MIC concentrations of polymyxin B and triclosan on the colistin-resistant *K. pneumoniae* strain. As shown in Figure 7, propidium iodide (PI) staining (red dots), marking dead cells, shows notable differences across treatments. The untreated control sample exhibits low cell death (∼11.2% PI+). Treatment with polymyxin B alone results in a modest increase in cell death to (∼27%), while triclosan alone leads to a slight decrease (∼7.4%), suggesting limited bactericidal efficacy at this sub-MIC level. However, the combination treatment significantly enhanced cell death (∼58.3%), suggesting a strong synergistic effect. The bar graph (Fig. 7), representing mean values from biological replicates with error bars, underscores this trend. Repeated measures ANOVA revealed a statistically significant difference among treatment groups (F(1.775, 3.549) = 106.7, p = 0.0007), with a high effect size (R² = 0.9816). Post hoc analysis indicates that the combination of polymyxin B and triclosan significantly increased cell death compared to either treatment alone or untreated controls. These findings suggest that sub-MIC combinations of these agents could effectively enhance bactericidal activity and potentially overcome AMR in colistin-resistant *K. pneumoniae*.

**Figure 7:**
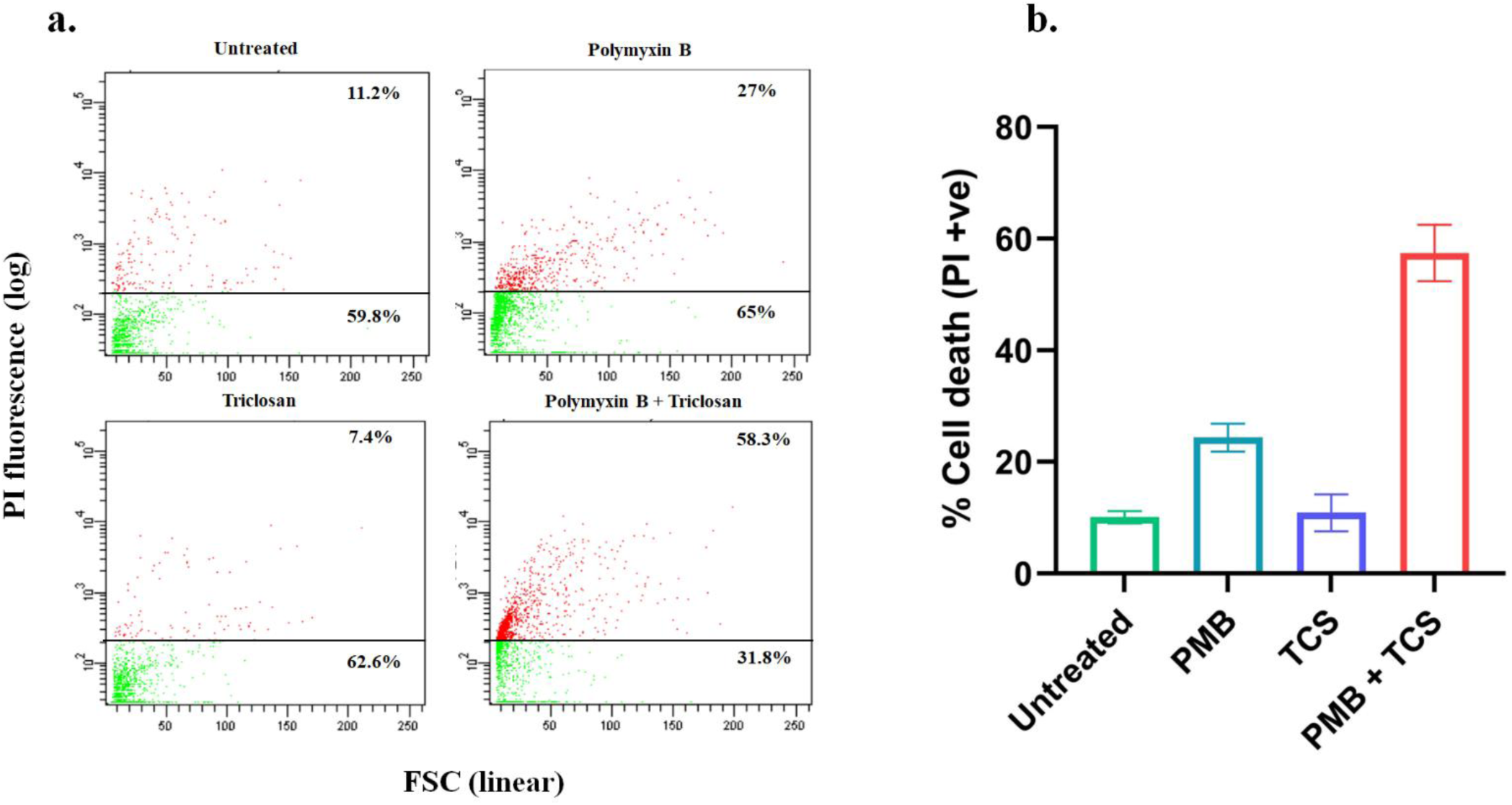
Flow cytometry analysis showing the effects of polymyxin and triclosan, individually and in combination, on cell death in a colistin-resistant *K. pneumoniae* (KPK1 strain). **(a.)** The four scatter plots represent untreated cells, cells treated with polymyxin B (2 µg/mL), cells treated with triclosan (0.0625 µg/mL), and cells treated with a combination of polymyxin B and triclosan at the same concentrations. Red dots indicate PI-positive (dead) cells, while green dots indicate live cells. Percentages in each plot represent the proportion of PI-positive cells, with the untreated control showing 11.2% cell death, polymyxin B alone at 27%, triclosan alone at 7.4%, and the combination treatment showing a significantly higher cell death rate of 58.3%, indicating a reversal of colistin resistance. **(b.)** The accompanying bar graph quantifies these percentages, demonstrating the enhanced bactericidal effect of the combination compared to individual treatments. Error bars represent standard deviation from triplicate of the sample (KPK1), and statistical significance was determined using two-way ANOVA.

### 2.8. Combination exhibits negligible cytotoxicity in RAW 264.7 cells

The MTT assay results, as shown in the bar graph (Fig. 8), demonstrate that treatment with polymyxin B, triclosan, and their combination (PMB + triclosan) did not significantly impact cell viability in RAW 264.7 cells. Across all treatment conditions, including the control, cell viability remained above 90%, indicating minimal cytotoxicity. Specifically, the treatments with polymyxin B and triclosan, individually and in combination, did not differ significantly from the control group, as confirmed by two-way ANOVA statistical analysis. These findings suggest that polymyxin B and triclosan are non-toxic to RAW 264.7 cells under the tested conditions.

**Figure 8:**
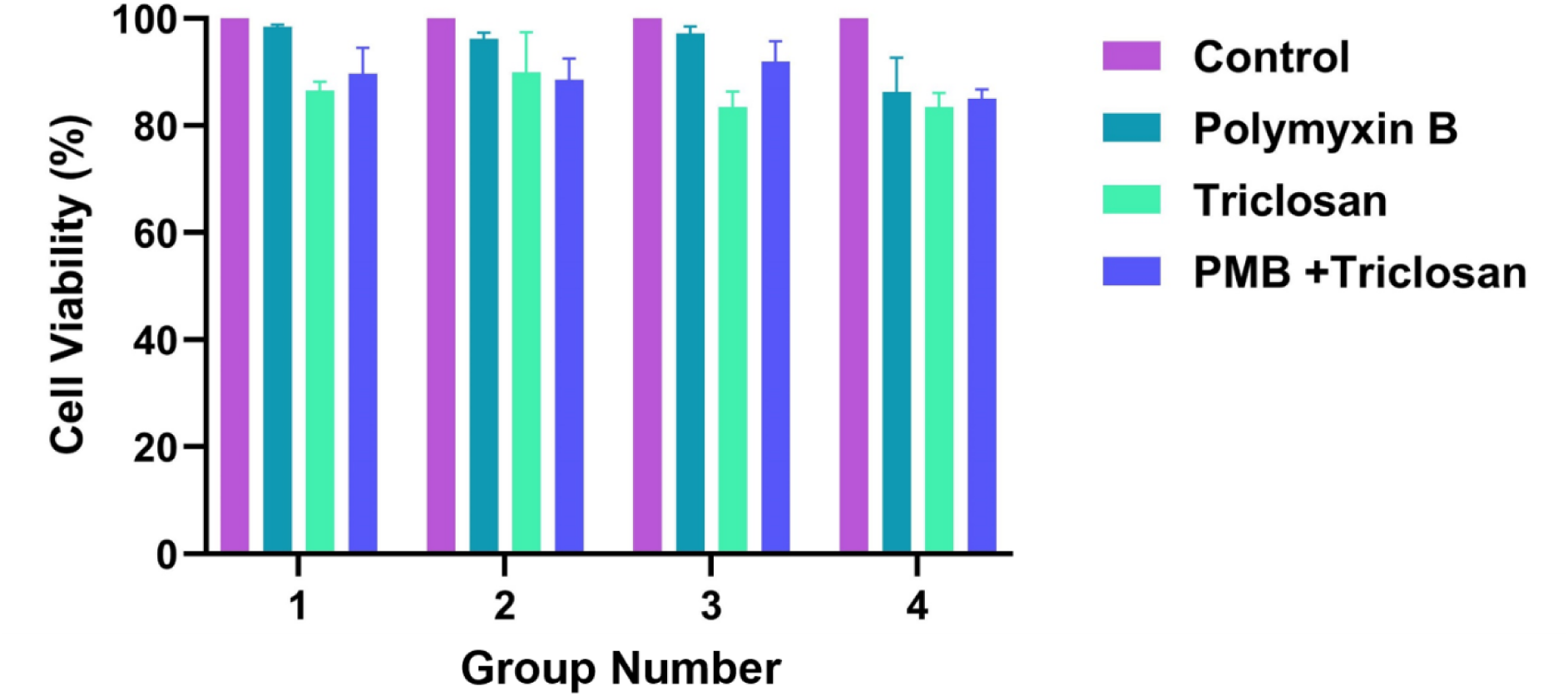
Cell viability of RAW 264.7 cells following treatment with varying concentrations of polymyxin B, triclosan, and their combination. Cell viability was assessed using an MTT assay. In Group 1, cells were treated with polymyxin B at 2 µg, triclosan at 0.031 µg, and a combination of 2 µg polymyxin B and 0.031 µg triclosan. In Group 2, concentrations were increased to 4 µg polymyxin B, 0.125 µg triclosan, and a combination of 4 µg polymyxin B and 0.125 µg triclosan. In Group 3, cells received 8 µg polymyxin B, 0.5 µg triclosan, and a combination of 8 µg polymyxin B with 0.5 µg triclosan. In Group 4, the highest concentrations were tested: 16 µg polymyxin, B1 µg triclosan, and a combination of 16 µg polymyxin B and 1 µg triclosan. Error bars represent standard deviation from triplicate samples, and statistical significance was analysed using two-way ANOVA.

**Figure 9:**
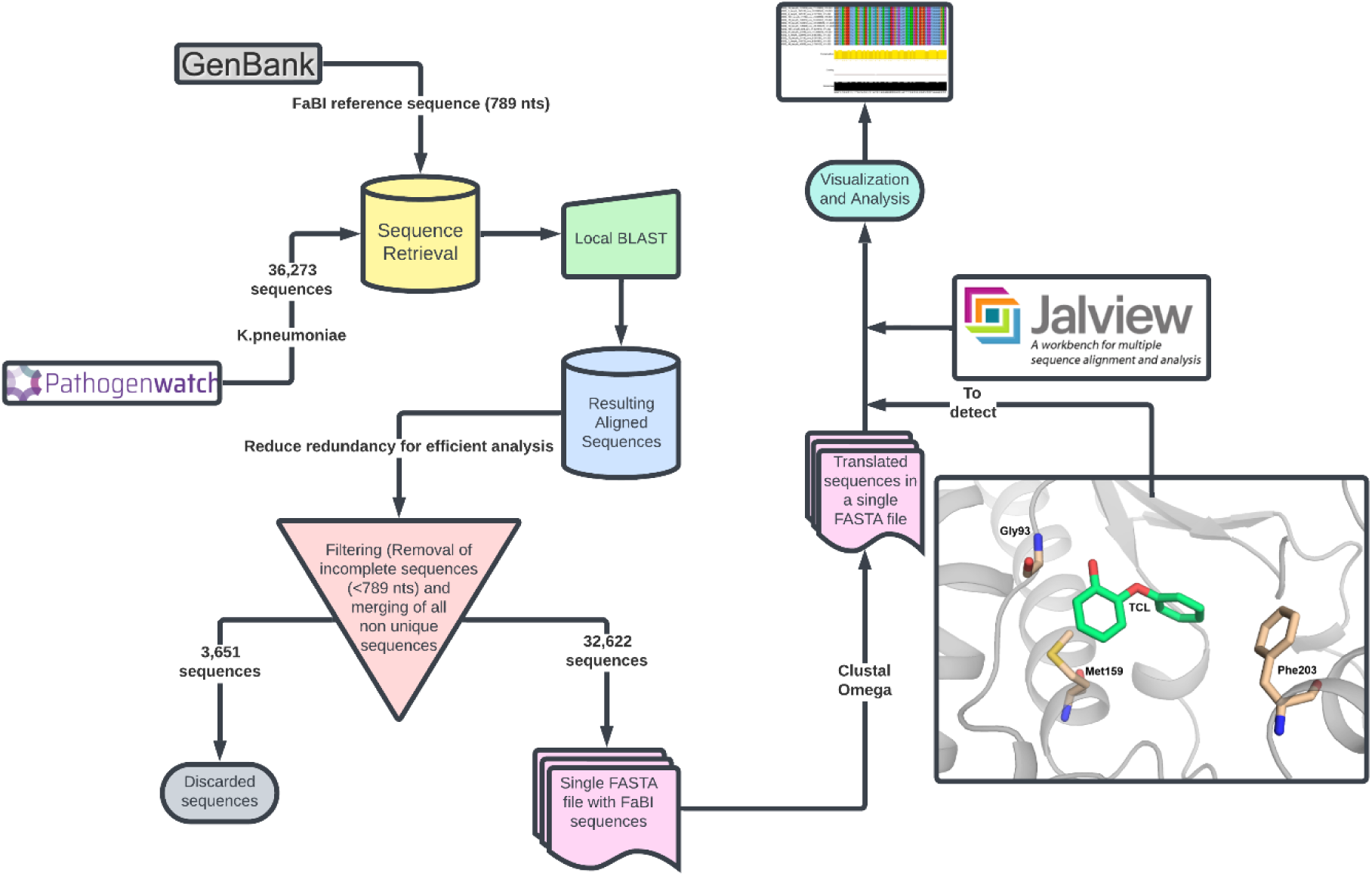
A schematic workflow for the detection of mutations in triclosan binding site residues of FabI, a target for triclosan binding. Sequences of FabI encoding gene were aligned and examined for mutations in the three binding site residues of FabI.

### 2.9. *Klebsiella pneumoniae* isolates do not have a significant mutation in *fabI* gene

Since several reports have hypothesized that resistance against triclosan occurs as it is released in the environment after its applications. Thus, we attempted to examine the potential mutation in the FabI encoding gene, as FabI is an exclusive target for triclosan activity. There are three potential mutations reported in the active site of FabI that may reduce the affinity of triclosan and subsequently confer resistance to triclosan. Among these reported residues, Gly93 has direct interaction through hydrogen bonds, while Met159 and Phe203 interact through hydrophobic and aromatic ring stacking attractions. Therefore, we focused on identifying mutations in these residues in *K. pneumoniae* isolates. Surprisingly, we did not find significant mutation of these three residues in isolates.

## 3. Discussion

Gram-negative bacterial pathogens with multidrug resistance (MDR) pose a significant global threat (Breijyeh et al., 2020), specifically, *K. pneumoniae* emerging as a leading cause of antibiotic resistance propagation (Alcántar-Curiel et al., 2018). In the context of AMR, *K. pneumoniae* strains producing extended-spectrum beta-lactamases (ESBLs) and showing carbapenem resistance have emerged as a significant concern, exhibiting resistance to the majority of clinically effective antibiotics (Karampatakis et al., 2023; Romyasamit et al., 2024). To address this crisis, polymyxin B and colistin have been extensively utilized as last-resort therapies for critically ill patients, particularly in ICU patients (Morris and Cerceo, 2020; Petrosillo et al., 2019). Alarmingly, resistance to colistin has also emerged, further exacerbating the threat and underscoring the urgent need for vigilant monitoring of its prevalence and underlying mechanisms (Goel et al., 2014).

The global prevalence of colistin resistance in *K. pneumoniae* from 2009 to 2023 reveals a rapid escalation in resistance rates during this period. By 2022, colistin-resistant *K. pneumoniae* accounted for over 13% of cases involving these infections. This alarming trend is exacerbated by the rise of plasmid-mediated mobile colistin resistance (*mcr*) genes, which facilitate the horizontal transfer of resistance across bacterial strains and species (Anyanwu et al., 2020). The first case of *mcr-1*-mediated colistin resistance in *K. pneumoniae* was reported in 2016 (Liu et al., 2016), and our findings corroborate this timeline, showing a sharp increase in *mcr-1* prevalence in subsequent years, peaking in 2023. Despite the significance of *mcr-1*, our data indicate that the majority of colistin-resistant *K. pneumoniae* strains acquire resistance through alternative mechanisms. Previous studies identified mutations in the *mgrB* gene, which encodes a transmembrane regulatory protein, as well as mutations in genes linked to the two-component regulatory systems PmrA/PmrB and PhoP/PhoQ, as major contributors to colistin resistance (Poirel et al., 2017; Wright et al., 2015). Enhanced expression of *pmrA, pmrB, pmrD, pmrK, phoP,* and *phoQ* genes has also been consistently observed in colistin-resistant isolates (Borsa et al., 2019). Furthermore, the continent-wise analysis of colistin resistance in *K. pneumoniae* revealed a marked global disparity in resistance patterns, underscoring the need for comprehensive antibiotic susceptibility profiling of clinical isolates.

In this time-elapsing situation, combination therapy has emerged as a swift strategy to combat MDR Gram-negative bacterial infections (Chang et al., 2022; Chen et al., 2024; Kaye et al., 2023; Lai et al., 2024; Li et al., 2024; Sirijatuphat et al., 2022). Clinically used combinations, such as beta-lactam antibiotics paired with aminoglycosides, have proven effective in treating severe infections caused by Gram-negative bacteria (Tamma et al., 2012). In the face of rising resistance to even last-resort drugs, combination therapies involving polymyxin B have shown great potential to enhance and preserve antimicrobial efficacy (Chung et al., 2016). In this series, the combination of polymyxin B and trimethoprim is already being used for treating eye infections, including acute bacterial conjunctivitis and blepharoconjunctivitis (Goodnough and Johnson, 1991; Rosenblatt and Stewart, 1974; Wagner, 1995). The pairing of polymyxin B with netropsin has demonstrated potent activity against MDR *Acinetobacter baumannii* (Chung et al., 2016), while its combination with minocycline has shown promise against carbapenem-resistant *K. pneumoniae* (Zhao et al., 2020). Notably, synergistic interactions have also been reported with tigecycline and avibactam-based β-lactam/β-lactamase inhibitor (BL-BLI) combinations (Li et al., 2022). These findings highlight the increasing therapeutic potential of polymyxin B, based combination strategies in enhancing antimicrobial efficacy against MDR bacterial strains.

Therefore, we assessed the synergistic efficacy of polymyxin B combined with the antibacterial agent triclosan. Remarkably, this combination reduced the MIC of polymyxin B to 1 µg/mL in colistin-resistant *K. pneumoniae* strains, effectively making them susceptible again. Although *mcr-1*–positive isolates showed a slightly reduced degree of synergy compared to *mcr-1*– negative strains, the combination still consistently achieved FICᵢ values within the synergistic range (FICᵢ < 0.5), supporting its potential against plasmid-mediated resistance. This outcome compares favourably with previous findings where polymyxin B showed synergy with non-antibiotic agents such as citalopram (FICᵢ ∼0.31), sertraline (FICᵢ ∼0.31–0.53), and spironolactone (FICᵢ ∼0.26), though many required supra-therapeutic concentrations (Otto et al., 2019). Triclosan, as a FabI inhibitor, showed strong synergy with colistin against *mcr-1*-expressing Enterobacteriaceae (FICᵢ ∼0.09–0.24) (Carfrae et al., 2023), while furanone C-30 also demonstrated synergistic activity (FICᵢ ∼0.04–0.5), though at comparatively higher concentrations (Zhang et al., 2021). In contrast, our triclosan–polymyxin B combination achieved robust synergy at low, sub-inhibitory concentrations, with FICᵢ values ranging from 0.0095 to 0.0925 in both *mcr-1* positive and negative *K. pneumoniae* strains, which is significantly lower than other FIC_i_ reported for combination with colistin and polymyxin B; demonstrating superior potency and broader applicability.

The time-kill assay (TKA) has been proven to be a crucial assay in preclinical modelling, providing detailed insights into the interaction between antimicrobial agents and bacteria (Bax et al., 2017; Filimonova et al., 2021). The results of TKA on the combination were highly encouraging: within 24 hours, the bacterial load was reduced by approximately 2 log_10_, and by 48 hours, complete bacterial eradication was observed. Comparatively, in recent studies combinations involving polymyxin B, such as with rifampin or minocycline, have demonstrated rapid bacterial reductions (3-4 log_10_ CFU/mL at 1 hour) against polymyxin B-resistant *K. pneumoniae*, though regrowth frequently occurred within 24-48 hours, and sustained killing was achieved only through triple-drug regimens requiring higher doses and complex combinations (Wang et al., 2024). Similar transient effects were observed with furanone C-30 combination, which produced >2 log_10_ reduction by 24 hours but required comparatively high concentrations (Zhang et al., 2021), as well as with other polymyxin-based combinations that often-necessitated additional agents for prolonged efficacy (Sopirala et al., 2010).

One major challenge in treating *K. pneumoniae* infections is its ability to form biofilms (Guerra et al., 2022). Remarkably, our combination demonstrated exceptional efficacy in the biofilm adherence assay at the same susceptible range of polymyxin B used in TKA. Likewise, in previous findings, antibiofilm synergy was observed with colistin-furanone C-30 and polymyxin B-resveratrol combinations against Gram-negative pathogens, including *P. aeruginosa* (Qi et al., 2022; Zhang et al., 2021). While both studies showed significant biofilm inhibition, they required higher compound concentrations, up to 50 mg/mL for furanone and 128 µg/mL for resveratrol. In contrast, our combination achieved potent biofilm suppression at low, subinhibitory concentrations, highlighting its potential as a more practical and broadly effective antibiofilm strategy. Another important factor in developing new antimicrobial agents is assessing their evolutionary potential to cause resistance in target organisms (Martinez and Baquero, 2000; Uddin et al., 2021). The frequency of resistance (FoR) assay helps determine the chance of resistant strains emerging in clinical settings (Evans and Titlow, 1998). In this study, polymyxin B alone at 8×MIC exhibited single-step mutation frequency. However, no mutations were observed when using the combination of polymyxin B and triclosan in the MIC range, indicating a significantly reduced risk of resistance development. Non-significant cytotoxicity, observed in RAW 264.7 cells, adds further evidence in favour of the combination, suggesting that it maintains cellular viability during treatment. Additionally, enhanced bacterial cell death observed in the flow cytometry-based cell viability assay further corroborates our findings, highlighting the efficacy of the combination in promoting bacterial eradication while maintaining minimal toxicity to cells.

Triclosan is widely used in several personal products as an antibacterial agent; thus, there are various *in vitro* studies have reported that it can cause resistance by mutating binding site residues in its target FabI enzyme (Heath et al., 1999; Sivaraman et al., 2003). Therefore, we examined publicly available genomes of *K. pneumoniae* isolates to detect these mutations. Remarkably, we could not find any significant mutation for potential binding site residues in the *K. pneumoniae* FabI enzyme. It suggests that triclosan may not cause mutations in FabI, conferring triclosan resistance in *K. pneumoniae.* Furthermore, our frequency of resistance assay for triclosan also indicates that triclosan resistance was not developed by applying a subinhibitory concentration of triclosan. These findings show that triclosan in combination with polymyxin B may not cause mutation in FabI for resistant development.

Collectively, our findings underscore the promising potential of the combination of polymyxin B and triclosan as a novel therapeutic approach, suggesting the need for further *in vivo* validation using animal studies.

## 5. Materials and methods

### 5.1. Global colistin resistance analysis

A total of 40,445 *Klebsiella pneumoniae* whole-genome sequences were downloaded from the PathogenWatch database (Argimón et al., 2021) spanning the years 2009 to 2023. From this dataset, sequences were systematically filtered to identify isolates annotated with colistin resistance. The filtering process focused on sequences marked as colistin-resistant, identified by examining the antibiotic resistance profiles for each sequence within the PathogenWatch platform. From these resistant sequences, we screened for the *mcr-1* gene, which is known to contribute to colistin resistance (Granata and Petrosillo, 2017). To confirm the presence of the *mcr-1* gene, we utilized the Basic Local Alignment Search Tool (BLAST) with optimized parameters to enhance detection sensitivity (Altschul et al., 1990). Specifically, we set a word size of 11 to improve the tool’s capacity to identify short or subtle sequence alignments, which are characteristic of resistance genes like *mcr-1*. We applied an E-value cutoff of 1e-5 to exclude lower-confidence alignments, thereby retaining only highly reliable matches. Additionally, we imposed a minimum percent identity threshold of 90%, focusing our analysis on sequences with high similarity to known *mcr-1* variants. To manage the impact of insertions or deletions-common within mobile genetic elements, we implemented gap penalties of -2 for gap opening and -1 for gap extension. The BLOSUM62 scoring matrix (Henikoff and Henikoff, 1992) was employed to optimize the detection of conserved regions within the *mcr-1* gene. Furthermore, low-complexity regions were excluded from the analysis to prevent misleading alignments. Only alignments covering 70–80% or more of the query length were deemed significant, ensuring a high-confidence dataset and robust identification of *mcr-1* sequences.

Following the identification of *mcr-1* positive sequences, all data were organized on a yearly basis from 2009 to 2023. This included individual cases reported each year, as well as the total number of colistin-resistant strains identified annually. Using these data, the percentage of colistin-resistant infections for each year was calculated. Additionally, we identified the unique cases with only *mcr-1* positive sequences and determined the proportion of colistin resistance attributable specifically to the presence of the *mcr-1* gene in *K. pneumoniae* isolates. All data were visualized and analysed using GraphPad Prism (version 9.5.1) to facilitate graphical representation and comparison between the number of colistin-resistant cases and the prevalence of *mcr-1* in the positive isolates across the study period.

### 5.2. Geographic distribution of colistin resistance

*K. pneumoniae* sequences exhibiting colistin resistance, previously retrieved from the Pathogenwatch database for the period 2009 to 2023, were analysed to determine the geographic distribution of resistance. Along with the sequences, all available metadata were downloaded, providing detailed geographic information regarding the origin of each sequence. This allowed for classification by continent, including North America, Europe, Asia, Africa, South America, and Australia. To obtain continent-based data, colistin-resistant sequences were aggregated and grouped according to their continent of origin. Although some metadata was incomplete, the analysis was conducted based on the available data. The data were then summarized with case counts sorted by continent to generate cumulative totals for the period 2009 to 2023. These results were visualized on a world map (created using Canva), with each continent highlighted according to the number of colistin-resistant cases reported.

### 5.3. Strains and culture

Six colistin-resistant *K. pneumoniae* were selected from earlier isolated strains from patient blood samples, with due approval from the concerned authority. Initially, an antibiotic susceptibility test was carried out using VITEK system (BioMérieux, France) to identify colistin-resistant strains of *K. pneumoniae*. As per EUCAST guidelines, broth microdilution assays were further performed to confirm colistin-resistant strains. Next, *mcr-1* mediated the colistin resistance was identified using PCR by amplification of *mcr-1* gene. Inoculum optimization for *K. pneumoniae* was performed using the laboratory strain *K. pneumoniae subsp. pneumoniae* (MTCC109) as a standard. Primary cultures of this strain were prepared in triplicate and incubated overnight with shaking at 150 rpm at 37°C in Mueller-Hinton Broth (MHB). After incubation, cultures were diluted with fresh MHB to achieve a range of optical densities (OD) at 600 nm, spanning from 0.1 to 1.0. Samples from each culture, adjusted to these specific OD values, were subjected to serial dilutions (10^-4^ to 10^-10^), with higher OD cultures undergoing greater dilution to ensure countable colony-forming units (CFUs) on agar plates. Using the drop plate method, 5 µl aliquots from each dilution were plated onto MHB agar in triplicate and incubated overnight to allow colony formation. CFUs were enumerated the following day, and the data were plotted to establish a correlation between CFU counts and their respective OD values (as shown in Fig. S1). This approach provides accurate CFU estimation based on OD, supporting optimal inoculum preparation for experimental protocols involving *K. pneumoniae*.

### 5.4. Determination of minimum inhibitory concentration (MIC)

Resistance to polymyxin B and MIC was assessed using the broth microdilution (BMD) method. This method, recommended by ISO 20776-1 (ISO 20776-1:2019) and EUCAST (Giske et al., 2022). A working stock solution of polymyxin B was prepared at a 2-fold higher concentration (64 μg/mL) in freshly autoclaved MHB. The assay was conducted in a 96-well plate, with 100 μL of MHB added to each well of a row, except for the first. In the first well, 200 μL of the highest polymyxin B concentration (64 μg/mL) was added. Serial 2-fold dilutions were then carried out by transferring 100 μL from the first well to the next, continuing until 2-fold dilution of the desired concentration range of 0.5 μg/mL to 32 μg/mL was achieved. The last 100 μL from the final well was discarded, and a control well without the antibiotic was included to check for normal bacterial growth. Overnight cultures of *K. pneumoniae* (clinical strains), adjusted to an optical density (OD) of 0.1 (corresponding to ∼10^8^ CFU/mL and previously optimized through bacterial growth estimation methods), were further diluted 100-fold to reach a bacterial concentration of 10^6^ CFU/mL. A 100 μL inoculum of this bacterial suspension was added to each well, resulting in a final volume of 200 μL per well. The 2-fold concentration of the drugs in MHB was appropriately diluted to achieve the desired experimental concentrations, while also maintaining the inoculation density at 5 × 10^5^ CFU/mL, a standard inoculation amount for performing MIC experiments.

### 5.5. Checkerboard assay

The synergy between the combination drug polymyxin B and triclosan was evaluated using the checkerboard assay method in a 96-well plate. Initially, a polymyxin B antibiotic working stock was prepared at concentrations 2-fold and 4-fold higher than the desired concentration in MHB. Here, polymyxin B was tested at desired concentrations of 32 μg and 64 μg; for 32 μg, working stocks of 64 μg and 128 μg were prepared, while for 64 μg, a 256 μg working stock was used. Freshly autoclaved MHB was dispensed at 100 μL into each well, excluding row A. Subsequently, 200 μL of the 2-fold working stock of polymyxin B was added to wells 1 to 11 in row A, with the 4-fold working stock added to the 12^th^ well. This arrangement ensured that the subsequent addition of another drug in the 12^th^ column would result in a dilution from 4-fold to 2-fold, maintaining symmetry along the row for the concentration of polymyxin B. Serial dilutions were then carried out from rows B to G, with an excess 100 μL discarded from row G to maintain a final volume of 100 μL in each well (excluding row H), thus achieving a wide concentration range of polymyxin B from 0.5 μg to 32 μg in the final 96-well plate following inoculation (as shown in Fig. S2).

In the subsequent step, the second drug, triclosan, was prepared as a main stock solution DMSO, with a subsequent sub-stock or working stock prepared in MHB at a 4-fold higher concentration than the highest desired range of triclosan. Here, triclosan was tested at desired concentrations of 0.5 μg and 2 μg, using working stocks of 2 μg and 8 μg, respectively. Then, 100 μL of the working stock of triclosan was added to the 12^th^ column, already containing 100 μL of serially diluted polymyxin B, resulting in a total volume of 200 μL and halving the concentrations of triclosan and polymyxin B from 4-fold to 2-fold higher. Serial dilutions of triclosan were performed from the 12^th^ column to the 2^nd^ column, with excess 100 μL of media discarded from the 2^nd^ column to ensure a uniform volume of 100 μL throughout the plate, thus creating a wide concentration range of triclosan from 0.0004 μg to 0.5 μg in the final plate. This process facilitated the determination of the MIC for each drug as well as for the combination. For instance, the 1^st^ column contained only polymyxin B, while row H contained only triclosan, and the well in row H-1 served as a growth control with no drug present (as shown in Fig. S2).

The overnight growth culture of six (3 *mcr-1* + and 3 *mcr-1* -) clinical strains of *K. pneumoniae*, known to be polymyxin B and colistin-resistant, was utilized for inoculation. The culture was diluted according to the pre-described method to achieve a final inoculum of 10 × 10^5^ CFU/mL. Subsequently, 100 μL of the inoculum was added to each well to achieve a final volume of 200 μL and a final bacterial inoculum of 5 × 10^5^ CFU/mL. Additionally, it is important to note that with the addition of the inoculum, the drug concentrations in each well were halved, aligning with the desired concentration range for the assay. This step ensured the maintenance of consistent experimental conditions and facilitated the evaluation of antimicrobial efficacy across the intended concentration spectrum. The 96-well plate was subsequently incubated for 20-24 hours at 37 °C to observe the final outcomes. After the incubation period, OD values were collected at 600 nm and plotted as a heatmap to facilitate better understanding and calculation of the fractional inhibitory concentration (FIC) using the formula:

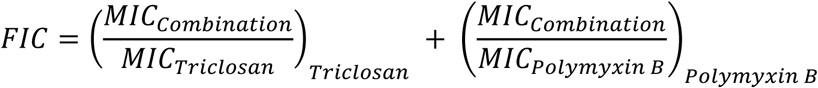

An FIC index of 0.5 or less indicates synergistic action between the antimicrobial agents, where their combined effect exceeds the sum of their individual effects. FIC indices falling between 0.5 and 4 suggest additivity, indicating that the combined effect is roughly equivalent to the sum of their individual effects. Conversely, an FIC index exceeding 4 indicates antagonism, signifying interference between the antimicrobial agents, leading to a diminished combined effect compared to their individual effects.

### 5.6. Synergy data analysis

OD data of KPK1 strain from the checkerboard assay were analysed using the Synergy Finder web application (Version 3.0) (Ianevski et al., 2022). Data were organized according to the reference model specified in the application, and due to software limitations, drug concentrations were converted from μg/mL to ng/mL. Drug combination responses were expressed as percent inhibition, calculated using the formula:

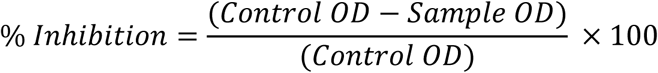

A four-parameter logistic regression (LL4) model was applied for curve fitting, and the Zero Interaction Potency (ZIP) model was used to assess synergy. Analysis results were then reported for interpretation.

### 5.7. Time-kill kinetics assay

The time-kill assay was conducted to evaluate the viability of *K. pneumoniae* (KPK1) against the combination of triclosan and polymyxin B. According to the Clinical and Laboratory Standards Institute (CLSI), the susceptible breakpoint MIC for polymyxin B and colistin in Enterobacterales is ≤2 μg/mL (CLSI; 2020). For this assay, the concentration of polymyxin B was set at 1/8 of the MIC, to get a final concentration of 2 μg/mL. Similarly, triclosan was also used at 1/8 of its MIC. Initial working stock solutions for both polymyxin B and triclosan were prepared at concentrations 200-fold higher than the desired assay concentrations, using freshly autoclaved MHB for polymyxin B and dimethyl sulfoxide (DMSO) for triclosan.

Freshly autoclaved tubes containing 2.5 mL of MHB were prepared for the assay. Four experimental conditions were set: one tube containing 1/8 MIC of polymyxin B, one containing 1/8 MIC of triclosan, one with a combination of both drugs and a control tube without any drug to allow for growth observation. To each tube, 25 μL of the respective working stock solution (polymyxin B or triclosan) was added to 2475 μL of MHB, achieving a 100-fold dilution for the individual treatments. For the combination, 25 μL of each working stock (polymyxin B and triclosan, totalling 50 μL) was added to the MHB, maintaining the same 100-fold dilution.

Subsequently, the *K. pneumoniae* bacterial inoculum was adjusted to 10^6^ CFU/mL using the previously described method, and an equivalent volume of 2.5 mL of the inoculum was added to each tube. This step ensured the attainment of the desired concentration of triclosan and a standardized bacterial inoculation count of 5 × 10^5^ CFU/mL. All tubes were then incubated at 37°C with 150 rpm. At predetermined time intervals of 1, 2, 4, 8, and 24 hours, samples were collected from each tube. Depending on the time elapsed and the turbidity of the tube, samples were serially diluted up to 10^-8^ to facilitate a readable CFU count on agar plates. Using the drop plate method, 5 µl aliquots from each serially diluted sample were plated onto MHB agar plates. For samples with higher concentrations of triclosan, aliquots were plated without dilution to accurately determine the rate of bacterial killing. All procedures, including triclosan dilution, bacterial inoculation, incubation, sampling, and CFU counting, were carried out independently in three sets of tubes or in triplicates to ensure experimental reproducibility and facilitate a comprehensive assessment of the efficacy of triclosan against *K. pneumoniae* over time and concentration.

All plates were incubated for 24 hours to enable proper CFU count determination. CFU counts for each concentration at each time point were calculated and converted into a logarithmic scale. The effectiveness of the combination over time and concentration was observed and recorded by plotting the value of log CFU against time points. All data analysis and visualization were performed using GraphPad Prism.

### 5.8. Adherent biofilm formation assay

Biofilm formation was quantified by measuring the absorbance at 595 nm (A595) of crystal violet-stained adherent cells, following a previously described method with slight modifications (Mandell et al., 2019). *K. pneumoniae* (KPK1) was initially cultured in 10 mL of LB broth and incubated overnight at 37°C. The overnight culture was then diluted to the optimal OD for inoculation, as determined by prior growth estimation protocols. Sub-inhibitory MIC concentrations of polymyxin B (2 μg/mL, equivalent to 1/8 MIC), triclosan (0.0625 μg/mL, also 1/8 MIC), and a combination of both were prepared in LB broth, resulting in a total volume of 100 μL per well. A 100 μL aliquot of bacterial suspension at 10^6^ CFU/mL was then added to each well, resulting in a 2-fold dilution and a final inoculum concentration of 5 × 10^5^ CFU/mL. Growth control wells and blank controls were included by adding 100 μL LB with 100 μL of bacterial culture and 200 μL of LB alone, respectively. The plates were incubated at 37°C for 24 hours (without agitation). Following incubation, the culture media was carefully removed from each well, and the wells were washed three times with distilled water to eliminate non-adherent or planktonic cells. Biofilms were stained by adding 200 μL of 0.01% crystal violet solution to each well, allowing the adherent cells to be properly stained. After 15 minutes, the excess stain was removed by washing twice with sterile water. Crystal violet retained by adherent biofilm cells was then solubilized using 30% glacial acetic acid, and absorbance at 595 nm was measured to quantify biofilm formation.

### 5.9. Frequency of resistance analysis

The spontaneous single-step mutation rates for *K. pneumoniae* (KPK1) were determined by growing bacteria in antibiotic-free MHB broth overnight (concentration of approximately 10^9^ CFU/mL) and plating 100 μL aliquots on Mueller-Hinton (MH) agar plate containing 4 x MIC, 8 x MIC of polymyxin B and triclosan individually, for the combination testing the MH agar plates were prepared in the MIC range (16 μg PMB + 0.5 μg TCS). A growth control was taken by serially diluting the overnight culture to 10^7^ to obtain countable colonies on MH agar plates, all plates were incubated for 24 to 48 hrs. All experiments were performed in triplicates. The resistance frequency was calculated using the following formula (Evans and Titlow, 1998):

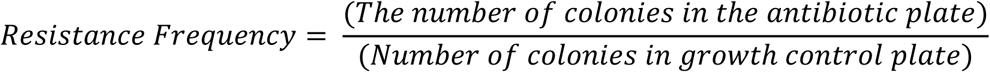

### 5.10. Cell viability assay

Clinical strains of *K. pneumoniae* were initially cultured overnight in MHB at 37°C. The following day, the bacterial suspension was adjusted to an optimized optical density corresponding to a final concentration of approximately 10^8^ CFU/mL, as previously determined. For the treatment phase, bacterial cells were exposed to sub-inhibitory concentrations of triclosan (0.0625 µg/mL, or 1/8 of the MIC) and polymyxin B (2 µg/mL, or 1/8 of the MIC). An additional combination treatment, maintaining the same concentrations of both agents, was prepared to assess potential synergistic effects. A control tube, containing the untreated bacterial culture, was included as a growth reference. All treatment and control tubes were incubated at 37°C for 4 hours.

Upon completion of the incubation period, 1 mL aliquots were withdrawn from each tube, including the control, and transferred into individual microcentrifuge tubes. To remove any residual antimicrobial agents, cells were pelleted by centrifugation at 7,000 *g* for 7 minutes, the supernatant was discarded, and the bacterial pellet was resuspended in 500 µl of sterile, syringe-filtered (0.22 µm) 1 X phosphate-buffered saline (PBS). A single wash step was performed by centrifuging the resuspended cells again at 7,000 *g* for 7 minutes, discarding the supernatant, and resuspending the pellet in fresh 1X PBS. To assess cell viability, the washed bacterial suspensions were stained using SYTO 9 (Thermofisher Scientific, USA) and propidium iodide (Thermofisher Scientific, USA) to distinguish live and dead cells. Staining was conducted by incubating the cell suspensions in the dark for 30 minutes. Following incubation, samples were analyzed using flow cytometry (BD FACS Canto II) to quantify the proportion of dead cells across each treatment condition, using event counts to assess bacterial viability.

### 5.11. MTT cytotoxicity assay

RAW 264.7 cells were cultured by seeding in a 96-well plate at a density of 10^4^ cells/well in 100 μL of complete cell culture medium, followed by incubation at 37 °C with 5% CO_2_ for 24 hours to allow cell attachment and recovery. After incubation, the cells were treated with a range of triclosan and polymyxin B concentrations to assess dose-dependent effects, as well as combination treatments of both agents. Specifically, four concentrations of triclosan were used: 0.031 μg, 0.125 μg, 0.5 μg, and 1 μg per well, along with four concentrations of polymyxin B: 2 μg, 4 μg, 8 μg, and 16 μg per well. For the combination treatments, cells received simultaneous doses of both compounds at various levels, including the highest concentrations of each agent: 16 μg of polymyxin B combined with 1 μg of triclosan. Additional combination treatments included 2 μg of polymyxin B with 0.031 μg of triclosan, 4 μg of polymyxin B with 0.125 μg of triclosan, and 8 μg of polymyxin B with 0.5 μg of triclosan.

Following the 24-hour treatment period, the medium in each well was carefully aspirated and replaced with 100 μL of fresh medium, followed by the addition of 10 μL of a 12 mM MTT stock solution (Himedia, India) prepared in sterile PBS. Cells were then incubated at 37°C with 5% CO_2_ for 3 hours to allow cellular reduction of MTT to formazan crystals. After this incubation, each well received 100 μL of a prepared solution of 0.01 M HCl in 10% SDS, which was added gently to dissolve the formazan crystals, facilitating measurement of the metabolic activity of the cells. The plates were then incubated for an additional 30 minutes to 1 hour at 37°C to ensure complete dissolution. Finally, each well was mixed by gentle pipetting, and the absorbance was measured at 570 nm to quantify the relative cell viability across the different treatment groups.

### 5.12. *fabI* gene variant analysis

Whole genome sequences of *K. pneumoniae* were obtained from PathogenWatch, a public genomic sequence repository and database. A total of 36,273 samples were taken (16th December 2023). The reference DNA sequence of *fabI* gene was obtained from *Klebsiella pneumoniae subsp. pneumoniae* MGH 78578 (GenBank Accession number CP000647.1:1441037-1441825). Local BLAST was carried out using the reference *fabI* gene sequence as a query, against the 36,273 samples to obtain the *fabI* sequence of each sample. Sequences with missing nucleotide residues and multiple gaps were omitted with a focus on the complete sequence length. After the removal, we were left with 32,622 sequences. The remaining sequences were then extracted in a single FASTA format file followed by the translation of all the *fabI* gene sequences obtained in the previous step using a local run of Clustal Omega (Sievers and Higgins, 2018). The output sequences were then arranged in a single FASTA file. The resulting sequences were then viewed in JalView (Waterhouse et al., 2009), and the mutations were analyzed and compared against previously reported mutations from the *fabI* gene of microbes other than *K. pneumoniae*. For ease of view, colours were added based on the alignment score.

## Statistical analyses

All experimental results were expressed as mean ± standard deviation (SD) of at least three independent biological replicates. Statistical analyses were performed using GraphPad Prism (version 9.5.1) and Synergy Finder (version 3.0) software tools. In graphical representations, error bars indicate standard deviations. For comparison of means between treatment and control groups, appropriate statistical tests were applied, including unpaired two-tailed t-tests and one-way or two-way ANOVA with relevant post hoc analyses (e.g., Student–Newman–Keuls method or Geisser-Greenhouse correction in repeated measures). Statistical significance was set at p < 0.05. Levels of significance were represented as follows: p < 0.05 (), p < 0.01 (), and p < 0.001 (), where applicable.

## Materials availability statement

All materials created in this manuscript may be accessed by contacting the corresponding author under appropriate material transfer agreements.

## Acknowledgments

Soumya Biswas is supported by the Bayer MEDHA Fellowship Program of Bayer Foundation. Gajraj Singh Kushwaha acknowledges Indian Council of Medical Research (ICMR) for Research Associate Award number BMI/11(45)/2020. We acknowledge the infrastructure support available through “Promotion of University Research and Scientific Excellence (PURSE)” program (SR/PURSE/2023/184) at KIIT University.

## Author contributions

Conceptualization and supervision: G.S.K. and M.S.; S.B. performed culture assay, synergistic, time-kill assay, S.G. carried out Computational studies; S.P. performed Cell Viability and MTT Assay, P.M. performed biofilm and resistance of frequency assay; G.V. provided clinical strain, N.M and M.S provided resources; S.B., S.G., S.P., GSK wrote first draft of the manuscript; S.B., G.S.K., and M.S. finalized the manuscript with comments from all other authors.

