## Supplemental Figure S1 and S2 for "Assessing the efficacy of therapeutically promising combination of polymyxin B and triclosan against colistin-resistant *Klebsiella pneumoniae*"

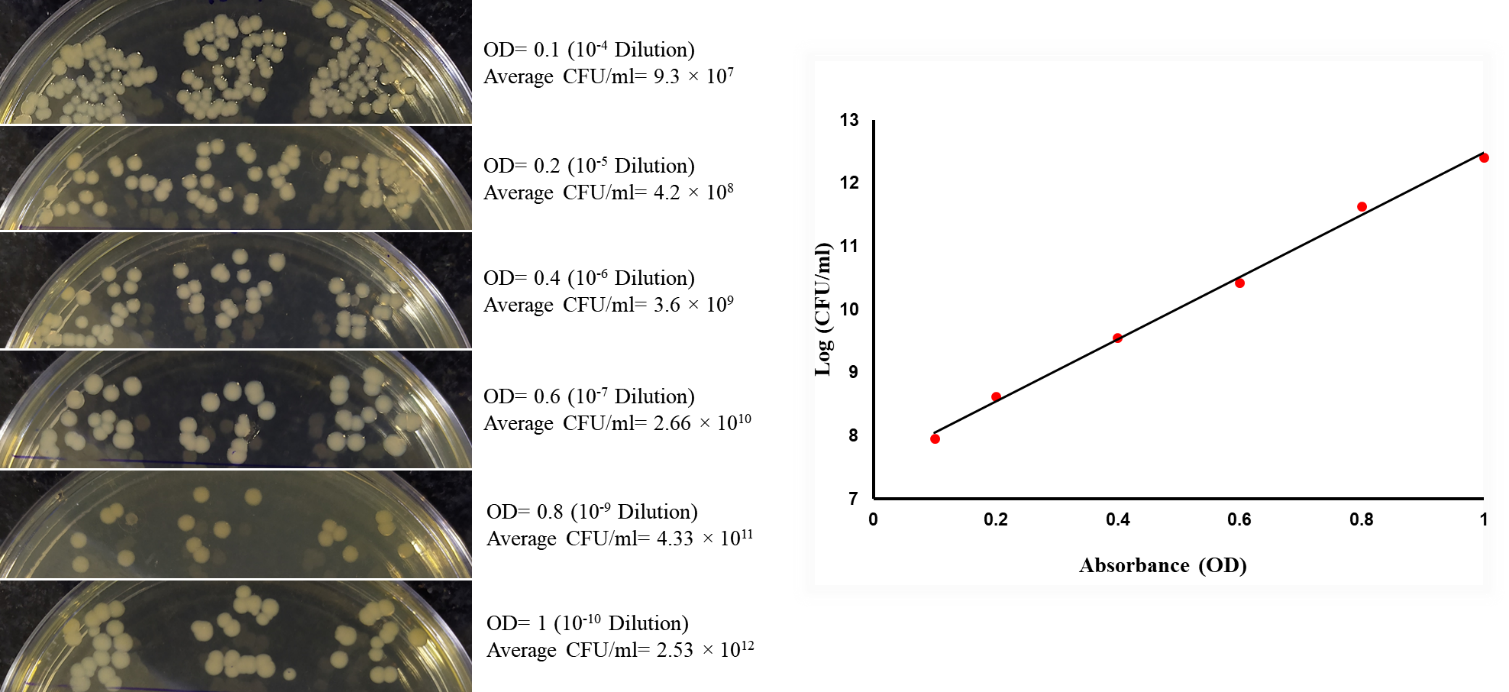

**Figure S1:** Inoculum optimization for *K. pneumoniae subsp. pneumoniae* (MTCC109). Cultures were grown in Mueller-Hinton Broth and adjusted to OD_600_ values ranging from 0.1 to 1.0. Serial dilutions (10⁻⁴ to 10⁻¹⁰) were plated using the drop plate method to determine colony-forming units (CFUs). CFU counts were correlated with corresponding OD values to standardize inoculum density.

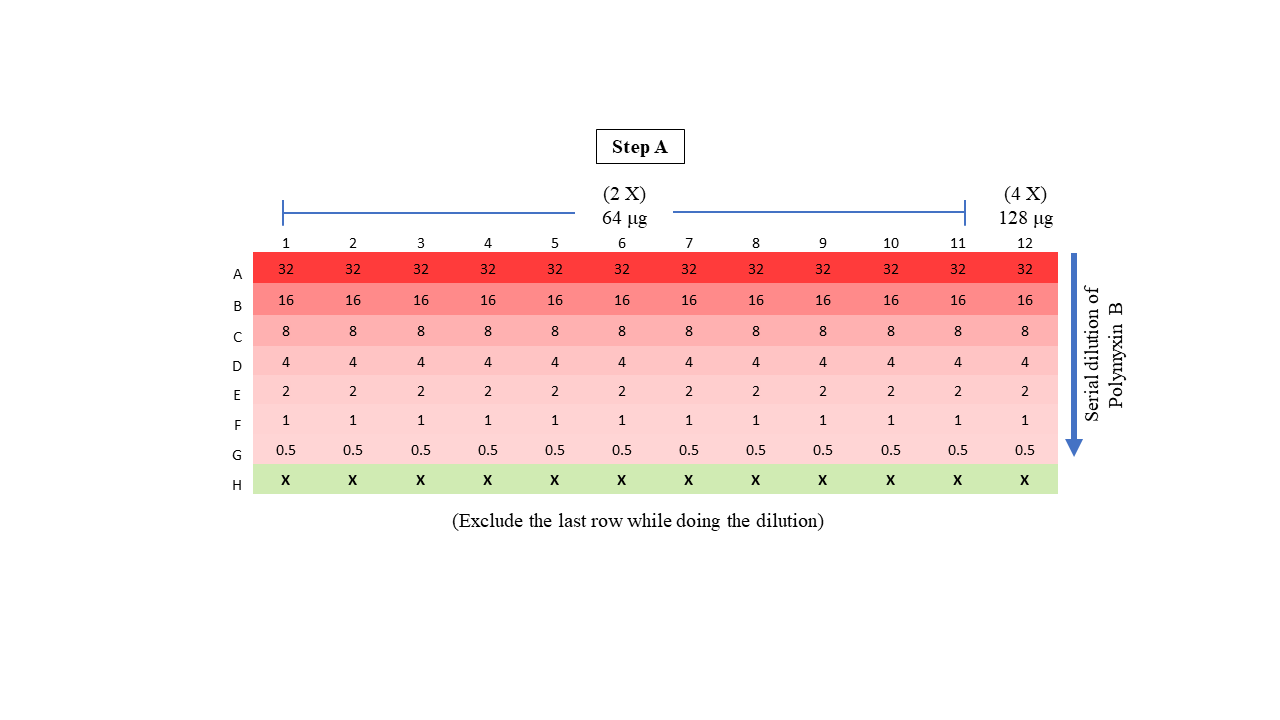

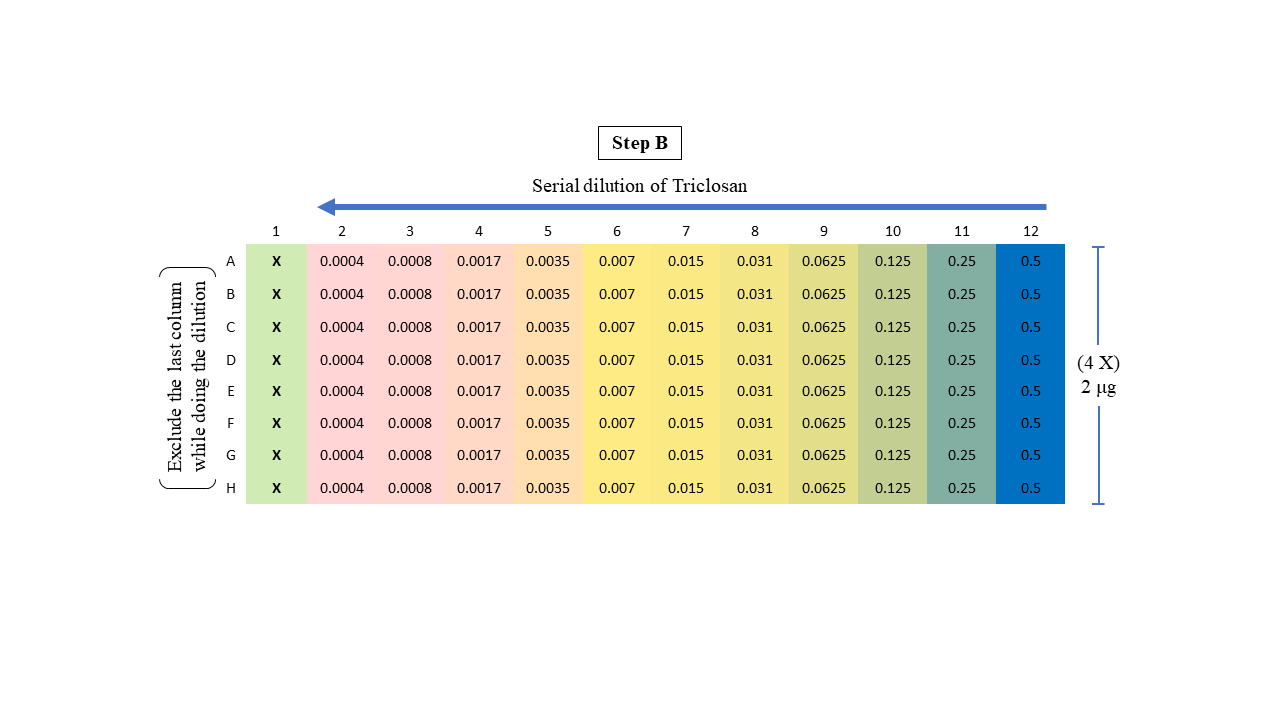

**Figure S2:** Checkerboard assay design for evaluating the combinatorial effect of Polymyxin B and Triclosan. Two-fold serial dilutions of Polymyxin B and Triclosan were prepared to achieve final concentration ranges of 32 to 0.5 µg/ml and 0.5 to 0.0004 µg/ml, respectively. Polymyxin B working stocks were prepared at concentrations of 64 µg/ml and 128 µg/ml to accommodate the desired dilution range. Triclosan stock solution was prepared at 2 µg/ml. Dilutions were performed in a 96-well microtiter plate in a checkerboard format, with Polymyxin B concentrations varying along the X-axis (columns) and Triclosan along the Y-axis (rows). Each well was inoculated with the bacterial suspension and incubated under standard conditions to assess synergistic or antagonistic interactions.
